# The DNA-relaxation-dependent Off-to-On biasing of the type 1 fimbrial genetic switch requires the Fis nucleoid-associated protein

**DOI:** 10.1101/2022.09.20.508701

**Authors:** Colin Conway, Michael C Beckett, Charles J Dorman

**Author notes:** Technical University of the Atlantic, Galway, Ireland (Colin Conway).

## Abstract

The structural genes expressing type 1 fimbriae in *Escherichia coli* alternate between expressed (phase ON) and non-expressed (phase OFF) states due to inversion of the 314-bp *fimS* genetic switch. The FimB tyrosine integrase inverts *fimS* by site-specific recombination, alternately connecting and disconnecting the *fim* operon, encoding the fimbrial subunit protein and its associated secretion and adhesin factors, to and from its transcriptional promoter within *fimS*. Site-specific recombination by the FimB recombinase becomes biased towards phase ON as DNA supercoiling is relaxed, a condition that occurs when bacteria approach the stationary phase of the growth cycle. This effect can be mimicked in exponential phase cultures by inhibiting the negative DNA supercoiling activity of DNA gyrase. We report that this bias towards phase ON depends on the presence of the Fis nucleoid-associated protein. We mapped the Fis binding to a site within the invertible *fimS* switch by DNase I footprinting. Disruption of this binding site by base substitution mutagenesis abolishes both Fis binding and the ability of the mutated switch to sustain its phase ON bias when DNA is relaxed, even in bacteria that produce the Fis protein. In addition, the Fis binding site overlaps one of the sites used by the Lrp protein, a known directionality determinant of *fimS* inversion that also contributes to phase ON bias. The Fis-Lrp relationship at *fimS* is reminiscent of that between Fis and Xis when promoting DNA-relaxation-dependent excision of bacteriophage λ from the *E. coli* chromosome. However, unlike the co-binding mechanism used by Fis and Xis at λ *attR*, the Fis-Lrp relationship at *fimS* involves competitive binding. We discuss these findings in the context of the link between *fimS* inversion biasing and the physiological state of the bacterium.

## INTRODUCTION

Type 1 fimbriae (or pili) are surface appendages found on members of the *Enterobacteriaceae* [Spaulding et al., 2018]. They are virulence factors in pathogenic strains [Connell et al., 1996; Flores-Mireles et al., 2015; Welch et al., 2002] and contribute to biofilm formation in the host [Anderson et al., 2003; Justice et al., 2004, Wright et al., 2007] and in the external environment [Xing and Marshall, 2020]. Fimbriae are up to 2 microns in length, similar to the length of an *Escherichia coli* cell, and 7 nanometres in diameter. Each of the fimbriae has a helical structure composed of repeated copies of the FimA subunit protein; the FimH adhesin is located at the tip where it is responsible for binding to mannose [Spaulding et al., 2018]. Mannose-sensitive agglutination of red blood cells is a diagnostic test for the presence of type 1 fimbriae [Abraham et al., 1985; McFarland et al., 2008; Spaulding et al., 2018].

The production of type 1 fimbriae is subject to phase variation, with fimbriate and afimbriate cells coexisting in the same population [Abraham et al., 1985; Eisenstein, 1981]. This behaviour has been interpreted as a bet-hedging strategy that balances the risks of producing these highly immunogenic fimbriae (detection by the host immune system; the physiological cost of making, exporting, and assembling the structures) with the benefits (biofilm-based community living; colonisation of a host or another environmental niche) [Jiang et al., 2019; Sanchez-Romero and Casadesus, 2021; Zamora et al., 2020]. The invertible *fimS* genetic element is the basis of phase-variable *fim* operon expression. This 314-bp DNA segment harbours both the promoter for the transcription of the *fim* operon (Fig. 1) and a Rho-dependent transcription terminator that influences the stability of the mRNA transcribed from the *fimE* gene [Hinde et al., 2005; Joyce and Dorman, 2002]. Inverting *fimS* connects/disconnects the *fim* operon to/from its transcription promoter, and connects/disconnects the *fimE* gene to/from its terminator, affecting FimE production [Hinde et al., 2005].

**Fig. 1.**
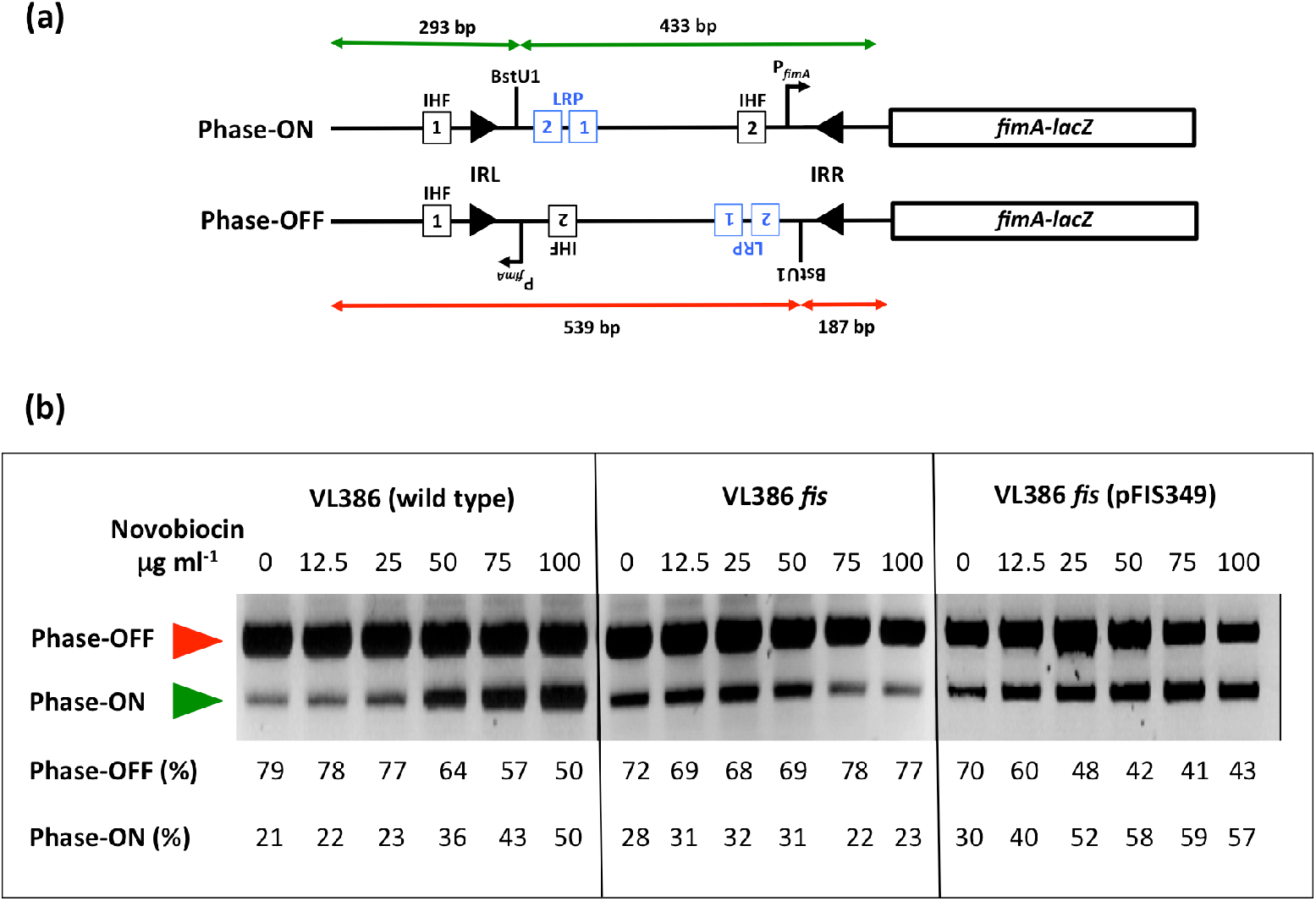
The Phase-OFF-to-Phase-ON bias of *fimS* genetic switch inversion in novobiocin-treated cultures is reversed in the absence of the Fis protein. (a) The *fimS* inversion assay. The *fimS* genetic element is amplified from the chromosome by PCR and the amplimers are cleaved with the BstUI restriction endonuclease (Methods). The lengths of the cleavage products are summarized for the ON (green) and OFF (red) orientations of *fimS*, above and below the drawings, respectively. The angled arrow, labelled P_*fimA*_, shows the position and orientation of the transcriptional promoter of the *fimA* gene. In strain VL386 and its derivatives, the *fimA* gene is fused to a *lacZ* reporter gene, allowing Phase-ON and Phase-OFF bacterial colonies to be distinguished on MacConkey lactose indicator agar plates. Squares represent the binding sites for IHF (black) and Lrp (blue), respectively. The filled arrowheads represent the left (IRL) and right (IRR) 9-bp inverted repeats that flank *fimS*. The position of the BstUI restriction endonuclease recognition site between Lrp binding site LRP-2 and IRR is shown. Not to scale. (b) Electrophoresis of the *fimS* DNA fragments from the wild-type strain (VL386), its *fis* knockout derivative, and the complemented *fis* mutant, following BstUI-digestion of the PCR-amplified *fimS* genetic element. The red arrowhead indicates the 539-bp Phase-OFF diagnostic band and the green arrowhead shows the 433-bp diagnostic band. The cultures had been treated with novobiocin at the concentrations given above each gel lane. The intensities of the DNA bands corresponding to the ON and OFF orientations of *fimS* in each lane were determined by densitometry and are reported as percentages below the lane. The experiment was performed three times and typical data are presented.

Inversion of *fimS* involves site-specific recombination within the 9-bp inverted repeats that flank the element [Gally et al., 1996; McCusker et al., 2008] (Fig. 1). In pathogenic strains of *E. coli*, the paralogous, independently acting, tyrosine integrases, FimB and FimE, promote inversion [Holden et al., 2007; Klemm, 1986; Kulasekara and Blomfield, 1999; Smith and Dorman, 1999]. These integrases catalyse the recombination reaction using the same chemistry, but their distinct DNA binding preferences at the alternate forms of *fimS* determine their recombination biases: FimB inverts both the phase ON and phase OFF forms of *fimS* with equal efficiency whereas FimE has a strong preference for the phase ON form, biasing FimE-mediated recombination in favour of ON-to-OFF switching [Gally et al., 1996; McClain et al., 1991; McCusker et al., 2008; Smith and Dorman, 1999]. In bacteria producing both FimB and FimE, the ON-to-OFF inversion preference of FimE predominates and inversion of the *fimS* element is biased strongly towards the OFF orientation [Gally et al., 1996; Hinde et al., 2005; Joyce and Dorman, 2002; McClain et al., 1991]. Many laboratory strains of *E. coli* K-12 lack the FimE recombinase due to mutations in the *fimE* gene; in these strains, *fimS* inversion depends on the unbiased FimB recombinase alone [Blomfield et al., 1991].

FimB-dependent inversion of *fimS* is sensitive to DNA supercoiling [Dove and Dorman, 1994; 1996; Muller et al., 2009], a feature that it shares with the Int tyrosine integrase recombinase of bacteriophage λ [Crisona et al., 1999; Landy, 2015]. Inhibition of type II topoisomerase activity by the drug novobiocin results in a dose-dependent relaxation of negatively supercoiled DNA and concomitant biasing of *fimS* inversion in favour of the ON orientation [Dove and Dorman, 1994]. In λ, DNA relaxation favours excision of the prophage from the chromosome while negative DNA supercoiling is required for efficient λ integration [Crisona et al., 1999; Landy, 2015]. Thus, in both of these tyrosine-integrase-mediated site-specific recombination systems, DNA topology exerts differential effects on the directionality of the reaction. Three nucleoid-associated proteins (NAPs) influence the inversion of *fimS*: the Integration Host factor, IHF [Blomfield et al., 1997; Dorman and Higgins, 1987; Eisenstein et al., 1987]; the Leucine-responsive Regulatory Protein, Lrp [Blomfield et al., 1993; Corcoran and Dorman, 2009; Gally et al., 1994; Kelly et al., 2006]; and the Histone-like Nucleoid Structuring protein, H-NS [Corcoran and Dorman, 2009; Higgins et al., 1988; Kawula and Orndorff, 1991; O’Gara and Dorman, 2000].

IHF is essential for inversion; in its absence the *fimS* element becomes frozen in either the ON or OFF orientation, reflecting the switch phase at the moment that IHF was removed from the cell [Dorman and Higgins 1987; Eisenstein et al., 1987]. IHF binds to two sites; the IHF-1 site is adjacent to the left inverted repeat (IRL) of *fimS* while site IHF-2 is within *fimS* (Fig. 1a). Site IHF-2 has an ancillary role in boosting the activity of the *fimA* promoter [Dorman and Higgins, 1987] while IHF-1 is essential for Phase-OFF-to-ON orientational bias [Corcoran and Dorman, 2009]. IHF acts in concert with Lrp to impose Phase-ON orientational bias; Lrp binds to two sites, LRP-1 and LRP-2 (Fig. 1), within *fimS* [Gally et al., 1994] and its presence is required for Phase-ON biasing when DNA is relaxed [Kelly et al., 2006]. Once Phase-ON bias is established, the H-NS NAP is required to maintain it. This is achieved when H-NS binds to *fimS* and to the adjacent chromosomal DNA, creating a nucleoprotein ‘trap’ that maintains *fimS* in the ON orientation under conditions of relaxed DNA topology [Corcoran and Dorman, 2009; O’Gara and Dorman, 2000]. Thus, Lrp acts as a directionality determinant in *fimS* site-specific recombination, by analogy with the role of the Xis protein during bacteriophage λ excision from the *E. coli* chromosome, catalysed by the Int tyrosine integrase [Papagiannis et al., 2007].

Lrp binds cooperatively to DNA [Chen et al., 2005] at sites matching a degenerate consensus sequence [Cui et al., 1999], its 8-mer/16-mer oligomeric structure is sensitive to L-leucine [Chen and Calvo, 2002] and its gene regulatory activities may be indifferent to, stimulated by, or inhibited by, L-leucine and other amino acids [Calvo and Matthews, 1994; Chen et al., 2001; Ziegler and Freddolino, 2021]. The production of Lrp is subject to transcriptional autorepression [McFarland and Dorman, 2008; Wang et al., 1994]; it has a broad impact on gene expression [Cho et al., 2008], and plays a central part in the adaptation of the bacterium to the stationary phase of the growth cycle [Kroner et al., 2019; Tani et al., 2002; Zinser et al., 2000].

In contrast to Lrp, the Factor for Inversion Stimulation, Fis, is produced predominantly in early exponential phase of growth [Ball et al., 1992; Bogue et al., 2020; O Cróinín and Dorman 2007; Osuna et al., 1995]. It influences a wide range of DNA transactions: site-specific recombination [Esposito et al., 2003; Papagiannis et al., 2007]; chromosome replication [Gille et al., 1991; Grimwade and Leonard, 2021; Inoue et al., 2016; Ryan et al., 2004]; transcription [Kelly et al., 2004; O Cróinín et al., 2006; Karambelkar et al., 2012]; and transposition [Bétermier et al., 1989; Weinreich and Reznikoff, 1992]. The Fis protein influences DNA topology at several levels: it regulates the transcription of *topA*, the gene that encodes DNA topoisomerase I [Cameron et al., 2011; Weinstein-Fisher and Altuvia, 2007]; it also regulates the expression of the *gyrA* and *gyrB* genes, encoding the alpha and beta subunits, respectively, of the heterotetrameric DNA gyrase [Keane and Dorman, 2003; Schneider et al., 1999]. In addition, Fis acts as a topological buffer to set local DNA topology by constraining plectonemically supercoiled DNA [Rochman et al., 2002; Muskhelishvili and Travers, 2003]. Negative DNA supercoiling stimulates the transcription of the negatively-autoregulated *fis* gene [Schneider et al., 2000].

Fis is required to maintain the OFF orientation of *fimS* in the presence of the FimE recombinase [Saldaña-Ahuactzi et al., 2022] suggesting that Fis is another directionality determinant affecting *fimS* site-specific recombination in the same direction as Lrp. Here, we explore the role of the Fis protein in FimB-mediated *fimS* inversion by monitoring the inversion preferences of this site-specific recombinase in the presence or absence of Fis and by identifying biochemically, and then disrupting genetically, a binding site for Fis within *fimS* that is essential for determining the inversion preference of FimB. This Fis binding site overlaps substantially LRP-2, one of the Lrp binding sites in *fimS*, a situation that is reminiscent of the overlapping binding sites used by Xis and Fis in the *attR* region of the λ prophage during Int-mediated excision of the bacteriophage from the chromosome [Landy, 2015; Papagiannis et al., 2007]. The Fis and Xis proteins bind simultaneously to *attR* [Papagiannis et al., 2007]; in contrast we found that Fis and Lrp bind competitively to the LRP-2 site in *fimS*.

## METHODS

### Media, growth conditions and genetic techniques

The strains used in these experiments were derivatives of *Escherichia coli* K-12 (Table 1). The VL386 Δ*fis*∷*kan* knockout mutant was derived by P1vir-mediated transduction [Miller, 1992; Sternberg and Maurer, 1991] using a CSH50 *fis*∷*kan* mutant lysate. VL386 *lrp*∷*cml* was also prepared by transduction, using a CSH50 *lrp*∷*cml* lysate. Complementation of the *fis* mutation was carried out using plasmid pFIS349, which is a single-copy plasmid based on the mini-F origin plasmid pZC320 [Wilson et al. 2001]. Bacteria were cultured in L Broth (made from Difco media components) or L agar (containing agar at 1.5% w/v). MacConkey-lactose agar plates [Miller, 1992] were used for Lac phenotype determination. Unless otherwise stated, liquid cultures were grown overnight at 37°C with aeration. Where appropriate, antibiotics were used at the following concentrations: carbenicillin (100 μg ml^−1^), chloramphenicol (25 μg ml^−1^) and kanamycin (20 μg ml^−1^). Plasmid DNA was introduced to bacterial cells by CaCl2 transformation [Sambrook and Russell, 2001] or electroporation using a Bio-Rad Gene Pulser as described in Hanahan *et al*. [1991].

**Table 1.** Bacterial strains and plasmids.

| Strain/plasmid | Relevant details | Reference/source |
| --- | --- | --- |
| **Strains** |  |  |
| VL386 | $\phi(fimA-lacZ)\lambda pL(209)fimE::IS1$ | Abraham et al. [1985] |
| VL386*recD* | VL386 *recD::Tn10* | Smith and Dorman [1999] |
| CJD2116 | VL386 $\Delta fis::kan$ | This work |
| CJD2117 | VL386 $\Delta lrp::cml$ | This work |
| CJD2119 | CJD2116 (pFIS349) | This work |
| VL386*fimS-dist* | VL386 with Fis/LRP-2 binding site disrupted | This work |
| XL-1 Blue | *recA1 endA1 gyrA96 thi-1 hsdR17 supE44 relA1 lac* [F' *proAB lacIqZ* M15 Tn10 (Tet<sup>r</sup>)] | Stratagene |
| **Plasmids** |  |  |
| pSGS501 | *fimB::kan*, *fimE::IS1*, $\phi(fimA-lacZ)$ cloned in pACYC184, switch phase OFF Cm<sup>r</sup> | Smith and Dorman [1999] |
| pFIS349 | pGS349 containing the *Salmonella enterica* serovar Typhimurium *dusB-fis* operon, Ap<sup>r</sup> | Wilson et al. [2001] |
| pLSB124 | pMAK705 containing wild type *fimB* and *sacB* gene from *Bacillus subtilis* | This work |
| pMAK705 | temperature-sensitive allele exchange vector | Blomfield et al. [1991b] |
| pMCL210 | cloning vector, p15A replicon | Nakano et al. [1995] |
| pMMC106 | pMCL210 cut with NheI and XbaI and religated to delete the *lac* promoter | McCusker et al. [2008] |
| pMMC108 | *fimS* cloned as a 550 bp fragment in the PstI site of pMMC106 | McCusker et al. [2008] |
| pSLD203 | *fimB* gene cloned in pUC18 | Dove and Dorman [1996] |
| pUC18 | ColE1 replicon, Ap<sup>r</sup> | Yanisch-Perron et al. [1985] |

### Molecular biological techniques

Plasmid DNA was isolated using Qiagen Midi columns or Wizard mini prep columns (Promega). Specific DNA fragments were isolated using an agarose gel extraction kit (Roche Applied Science). Restriction enzyme digests were carried out using enzymes purchased from New England Biolabs by following the manufacturer’s recommended procedure. Plasmid DNA was sequenced using a T7 sequencing system (USB). Plasmid pSGS501 was used as the template with oligonucleotide COL69 as the sequencing primer (Table 2). Automated sequencing was carried out at MWG Biotech. This company also synthesized the oligonucleotides used in this study.

**Table 2.**
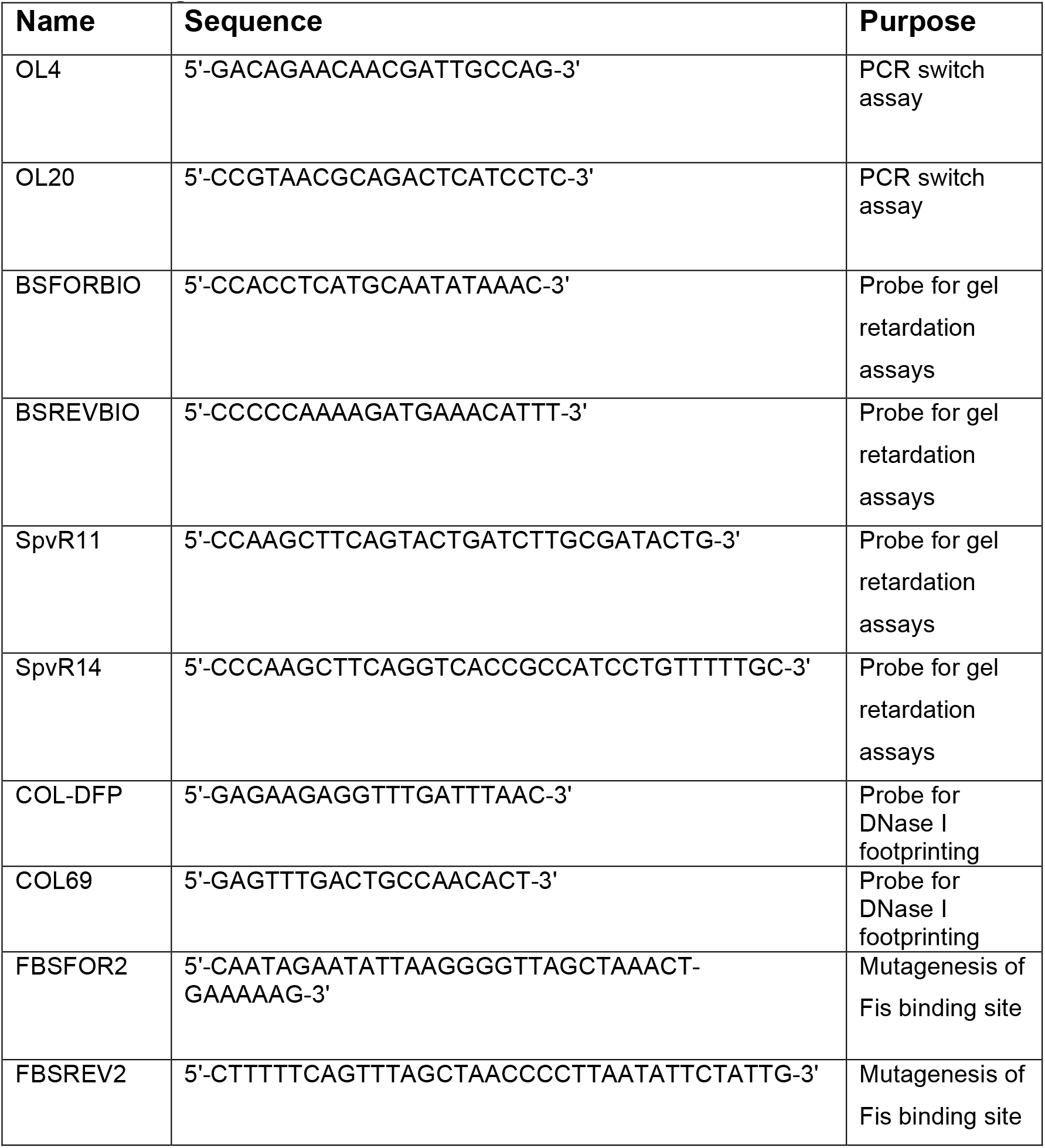
Oligonucleotides.

### Determination of *fimS* orientation on the chromosome

A PCR-based assay was used to determine the orientation of the *fimS* genetic switch on the *E. coli* chromosome (Fig. 1a). This method exploited the presence of a unique BstUI restriction site in the *fim* switch, *fimS*, which results in products of different lengths depending on the switch orientation [Smith and Dorman, 1999]. This restriction fragment length dimorphism allowed ON and OFF switches to be distinguished and quantified. Bacterial samples were harvested by boiling 50 μl of culture following overnight incubation at 37°C. Oligonucleotides OL4 and OL20 (Table 2) were used to amplify the switch region and generate a 726-bp DNA product. DNA amplification used Taq polymerase (New England Biolabs) with the following PCR conditions: denature at 94°C for 3 min, followed by 30 cycles of 94°C for 1 min, 58°C for 1 min and 72°C for 1 min. This was followed by a final extension time of 10 min at 72°C. Samples were cooled to 60°C, 10 units of BstUI were added to each reaction and incubation was continued at 60°C for 3 h. Digested PCR products were electrophoresed on 2% agarose gels. Phase OFF populations of bacteria yielded two DNA fragments of 539 and 187 bp in length, whereas phase ON populations gave fragments of 433 and 293 bp. The well-resolved 539 bp and 433 bp DNA fragments were used to commute the relative quantities of ON and OFF switches in the bacterial population: QUANTITY ONE image analysis software was used to measure approximate proportions of the resultant fragments (Fig. 1b).

### Analyzing protein binding to DNA by electrophoretic mobility shift assay

The association of purified Fis or Lrp proteins with the *E. coli fim* switch was measured using an electrophoretic mobility shift assay (EMSA). A 135-bp probe was amplified by PCR with Pfu polymerase (Stratagene), using the primer pair BSFORBIO and BSREVBIO (Table 2). The *S. enterica spvR* promoter was amplified using the primer pair, spvR11 and spvR14 and was used as a negative control for the Fis binding experiments. The probes were then purified using a PCR clean up kit (Roche Applied Science). The oligonucleotides had been ordered with 5’ biotinylated ends allowing for subsequent complex detection. Complexes were formed following incubation of amplified probe with increasing concentrations (0-270 nM) of purified His-tagged Fis for 15 min as described by the manufacturers of the Electrophoretic Mobility Shift Assay Kit (Pierce). Competitive binding of Fis and Lrp was tested in a similar manner with both proteins being added prior to DNA or binding buffer addition. Protein-DNA complexes were resolved by electrophoresis through a 7.5 % polyacrylamide gel for 2 h at room temperature. The gel was then electrophoretically blotted and developed using the procedure recommended by the manufacturer (Pierce).

### DNase I footprinting

A fragment encompassing *fimS* was amplified from pSGS501 with the primers COL69 and COL-DFP (Table 2). The PCR product was purified with a PCR clean-up kit (Roche Applied Science) and end-labelled with T4 polynucleotide kinase (New England Biolabs). This fragment was then digested for 2 h with MfeI at 37°C in a reaction volume of 60 μl. The probe was purified by extraction from a 6% polyacrylamide gel, following electrophoresis in TBE buffer. Labelled DNA was eluted in 3 ml of elution buffer [10mM TRIS-HCl pH 8.0, 1mM EDTA, 300 mM sodium acetate (pH 5.2), 0.2% SDS] at 37°C for 48 h. The eluted probe was extracted with an equal volume of phenol:chloroform and ethanol precipitated. The DNA pellet was then resuspended in 100 μl of double-distilled water. Two microliters of labelled probe solution were used in each footprinting experiment. DNA-protein complexes were formed in 50 μl of footprinting buffer (20 mM TRIS-HCl pH 7.5, 80 mM NaCl, 1 mM EDTA, 100 μg/ml BSA, 10% glycerol and 1 mM DTT) at 37°C for 30 min. Then 50 μl of 10mM MgCl2-5mM CaCl2 were added and incubation continued for a further 10 min. Next, 0.01 U of DNase I (Roche Molecular Biochemicals) was added, and digestion was allowed to proceed for 1 min. The reaction was terminated by the addition of 90 μl of stop solution (200 mM NaCl, 30mM EDTA pH 8.0, 1% SDS, 100 μg/ml tRNA). Samples were extracted once with an equal volume of phenol:chloroform, then precipitated with ethanol and resuspended in 6 μl of gel loading dye. Samples were denatured at 95°C for 3 min and were subjected to electrophoresis on a 7% urea-polyacrylamide gel alongside DNA sequencing reactions.

### Site-directed mutagenesis and allele replacement

Site-directed mutagenesis was performed using the Quikchange II (Stratagene) site-directed mutagenesis kit, according to the manufacturer’s recommendations. The oligonucleotides used to mutate the Fis binding site (FBSFOR2 and FBSREV2) are described in Table 2, and were supplied by MWG Biotech. Plasmid pMMC108 [McCusker et al., 2008] was used as the substrate for the mutagenesis. The method of allele replacement was as described in Smith and Dorman [1999]. Briefly, the mutated Fis binding site was introduced to the chromosome by cloning an MfeI-SnaBI fragment of *fimS*, containing the disrupted site, into pSGS501, a plasmid containing the *cat* chloramphenicol resistance gene (Table 1). The resulting plasmid was digested with EcoRV and an 8-kb fragment containing the mutated *fimS* region was gel extracted. Two μg of this fragment were electroporated into strain VL386*recD*. Loss of plasmid sequences following homologous recombination with the chromosome was confirmed by testing the transformants for chloramphenicol sensitivity. The presence of the disrupted Fis binding site in the chromosomal *fimS* element was confirmed by PCR amplification followed by DNA sequencing.

## RESULTS

### Loss of Fis alters the pattern of *fimS* inversion when DNA gyrase is inhibited

Inversion of *fimS* by the FimB recombinase becomes biased towards the ON orientation when the introduction of negative supercoils by DNA gyrase is inhibited by novobiocin. Increasing the dose of the gyrase-inhibiting drug exacerbates this effect. The Fis protein is known to preserve negative supercoils at a local level by binding to DNA [Rochman et al., 2002; Muskhelishvili and Travers, 2003]. We investigated the impact of eliminating Fis protein production on the inversion of *fimS* by FimB, using a wild type *E. coli* strain and an isogenic *fis* knockout mutant, treated with increasing concentrations of novobiocin (Fig. 1b).

In the wild type, treatment with incrementally increasing concentrations of novobiocin was accompanied by a progressive accumulation of *fimS* switches in the ON orientation, in agreement with previous data [Dove and Dorman, 1994]. In the *fis* mutant, this effect was not observed: as the concentration of the drug increased, the switch orientation remained close to a constant ratio of 30% ON and 70% OFF (Fig. 1b). The introduction of a functional copy of *fis* on a single copy plasmid restored the biased OFF-to-ON switching that is characteristic of the wild type (Fig. 1b). These data implicated Fis as a contributing factor in biasing FimB-mediated *fimS* switching when DNA gyrase activity is inhibited by novobiocin. We decided to investigate the relationship between Fis and *fimS* in more detail at the molecular level.

### Characterisation of a Fis binding site within *fimS*

Although the consensus sequence of the Fis binding site in DNA is degenerate, it has a number of features that are highly conserved among high-affinity sites [Hancock et al., 2016; Stella et al., 2010]. These allowed us to identify a potential binding site for Fis within the *fimS* element, located 50 bp from the 9-bp inverted repeat that forms the right-hand boundary of the switch when in the OFF orientation (Fig. 2). We used DNase I footprinting to map the binding site of purified *E. coli* Fis protein on the *fimS* genetic element *in vitro*. The site that was protected by Fis from DNase I digestion corresponded to the DNA sequence that matched with the consensus for high-affinity Fis binding sites (Fig. 2). The DNase I footprint consisted of bases that were protected by Fis and bases that became hypersensitive to digestion in the presence of the NAP. The latter are commonly found in sites occupied by DNA-binding proteins that bend DNA, a known property of Fis [Gille et al., 1991; Gosink et al., 1993; Hancock et al., 2016; Stella et al., 2010].

**Fig. 2.**
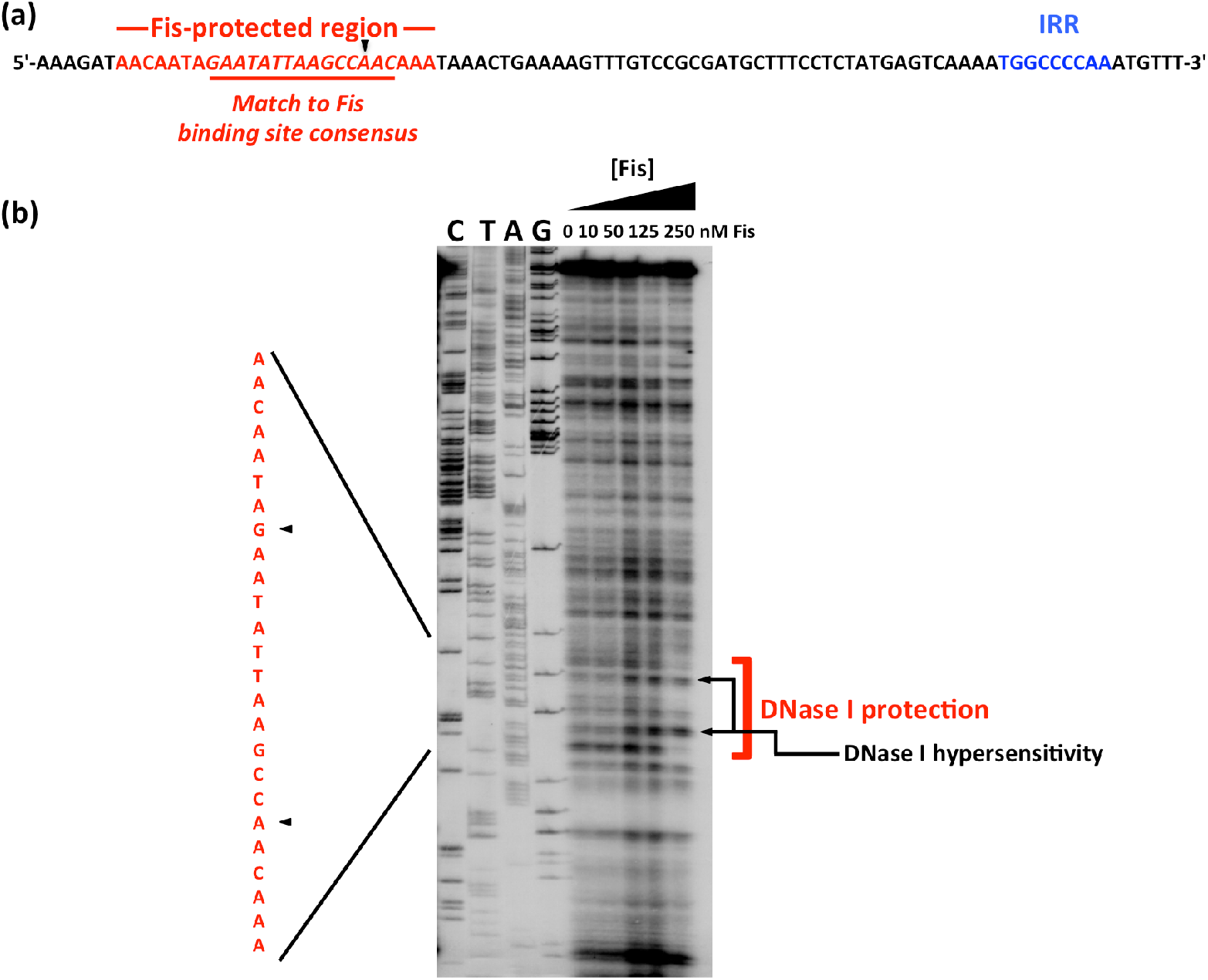
Identifying a binding site for Fis within *fimS*. (a) The DNA sequence of the right end of *fimS* in Phase-OFF, showing the location of the right inverted repeat, IRR (blue), and a sequence matching the consensus for Fis binding sites (red). (b) DNase I footprinting was performed with purified Fis protein and a DNA fragment corresponding to the right end of Phase-OFF *fimS*. Concentrations of Fis are given above each lane. The products of dideoxy chain-terminator nucleotide sequencing reactions, carried out with the same DNA fragment, are shown in the lanes labelled C, T, A, and G. The assay revealed a region in which Fis protected the DNA from DNase I digestion, with two nucleotides exhibiting hypersensitivity to the enzyme. The protected region is highlighted with a red bracket and black arrows indicate the two hypersensitive bases. Black arrowheads point to the regions of hypersensitivity in the sequence shown at left in red; this sequence corresponds to that shown in red in (a).

Further evidence of Fis binding to *fimS* came from an electrophoretic mobility shift assay (Fig. 3). At 90 nM Fis, the protein formed a complex that was consistent with the occupation of a single binding site. The *spvR* promoter region from *Salmonella enterica* serovar Typhimurium, that does not bind Fis [Kelly et al., 2004], was used as a negative control. This DNA sequence did not form a complex with Fis, at this or a higher concentration of the protein (Fig. 3).

**Fig. 3.**
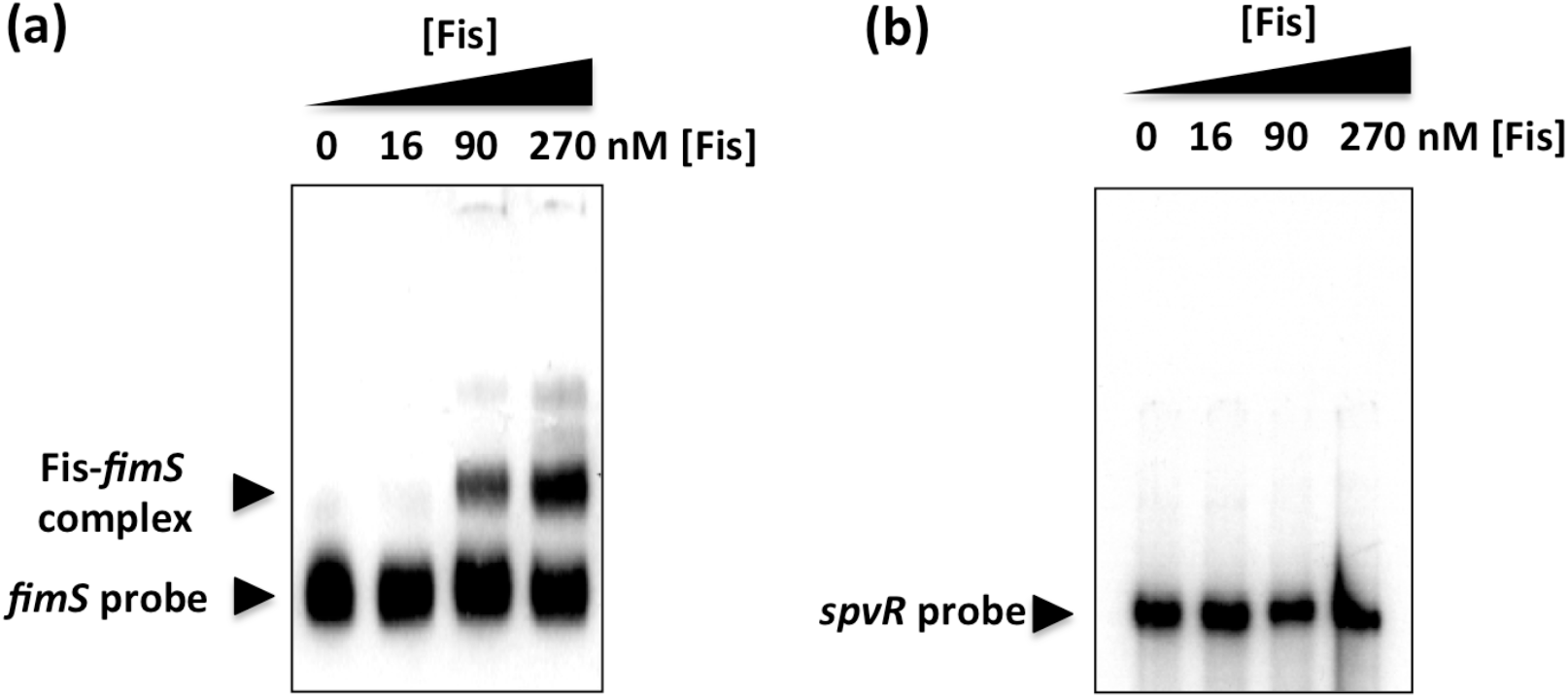
Demonstrating Fis binding to *fimS* by electrophoretic mobility shift assay. (a) Purified Fis protein was incubated at increasing concentrations with a labelled *fimS* fragment that corresponded to the *fimS* DNA sequence analysed by DNase I footprinting (Fig. 2b). Arrowheads indicate the positions of the protein-free probe and the Fis-*fimS* complexes on the gel. (b) A control EMSA using a DNA fragment corresponding to the *spvR* promoter region from *S*. Typhimurium. This DNA fragment was chosen because it is known not to bind Fis [Kelly et al., 2004]. Purified Fis failed to alter the electrophoretic mobility of the *spvR* DNA fragment at the same protein concentrations used in (a). These concentrations are shown above each gel lane in (a) and (b).

The Fis binding site within *fimS* was subjected to base substitution mutagenesis to alter its DNA sequence without altering its length. Changes were made to eight contiguous bases, destroying the match to the consensus sequence for high-affinity Fis binding sites (Fig. 4). The modified DNA element could not bind Fis at a protein concentration of 90 nM and only a weak interaction was detected at 270 nM (Fig. 4) that was likely due to the known tolerance of Fis for mismatches to its binding site consensus sequence (Hancock et al., 2016). Taken together, the DNase I footprinting data and the EMSA results show that Fis binds to the *fimS* genetic switch at a site that is located 50 bp from the inverted repeat boundary at the *fimA-*promoter-distal end of *fimS*.

**Fig. 4.**
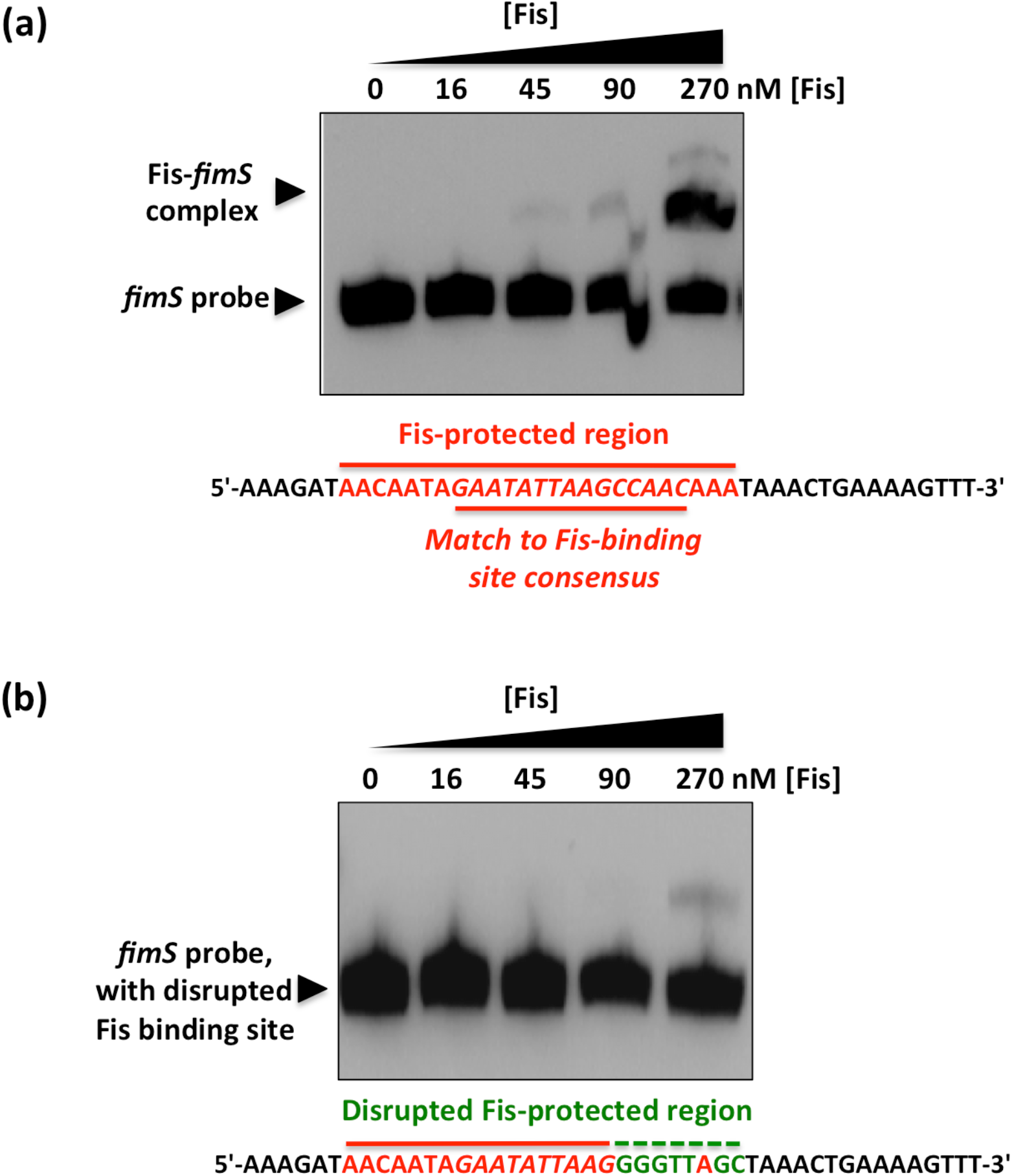
Mutation of the Fis binding site abrogates the Fis-mediated electrophoretic mobility shift of *fimS* DNA. (a) EMSA showing the gel mobility shift of a labelled fragment of *fimS* DNA that includes the Fis binding site, highlighted in red below the gel. (b) Disruption of the Fis binding site by substituting the bases, shown below the gel in green, of the wild type binding site sequence (red) almost completely abrogated the ability of Fis to alter the electrophoretic mobility of this DNA fragment. The concentrations of purified Fis used in the experiments are given above each gel lane; black arrowheads indicate the bands corresponding to the unbound *fimS* DNA probe and the Fis-*fimS* complex.

### The Fis binding site is crucial for *fimS* inversion preferences

The derivative of *fimS* with the 8-bp substitution mutation in the Fis binding site was transferred to the *E. coli* chromosome by homologous recombination. The mutant strain and the wild type were treated with increasing concentrations of novobiocin and the orientation of *fimS* was monitored by PCR (Fig. 5a). In the wild type, the *fimS* element adopted a dose-dependent preference for the Phase-ON orientation with novobiocin treatment; in the mutant with the disrupted Fis binding site, *fimS* adopted a novobiocin-dependent preference for the OFF orientation, the opposite to the situation seen in the wild type (Fig. 5a). Not only had the direction of the inversion bias been reversed compared to the wild type, the response to novobiocin occurred at the lowest concentration of the drug (12.5 μg/ml). These results demonstrated that the Fis binding site plays a pivotal role in determining both the direction of the DNA inversion response and the sensitivity of *fimS* inversion to DNA gyrase inhibition. Inspection of the Fis binding site’s location suggested that it overlapped Lrp binding site LRP-2 (see next section below) and the base substitution mutations might also have impaired Lrp binding to that site. For this reason, the strains used in the *fimS* orientation assay contained a plasmid pUC18 derivative, pSLD203, over-expressing the FimB recombinase (Table 1). This is an established way to allow *fimS* inversion to continue in strains deficient in co-factor production/binding without affecting the response of *fimS* recombination to DNA relaxation (Corcoran et al., 2009; Dove and Dorman, 1996; Kelly et al., 2006; Smith and Dorman, 1999).

**Fig. 5.**
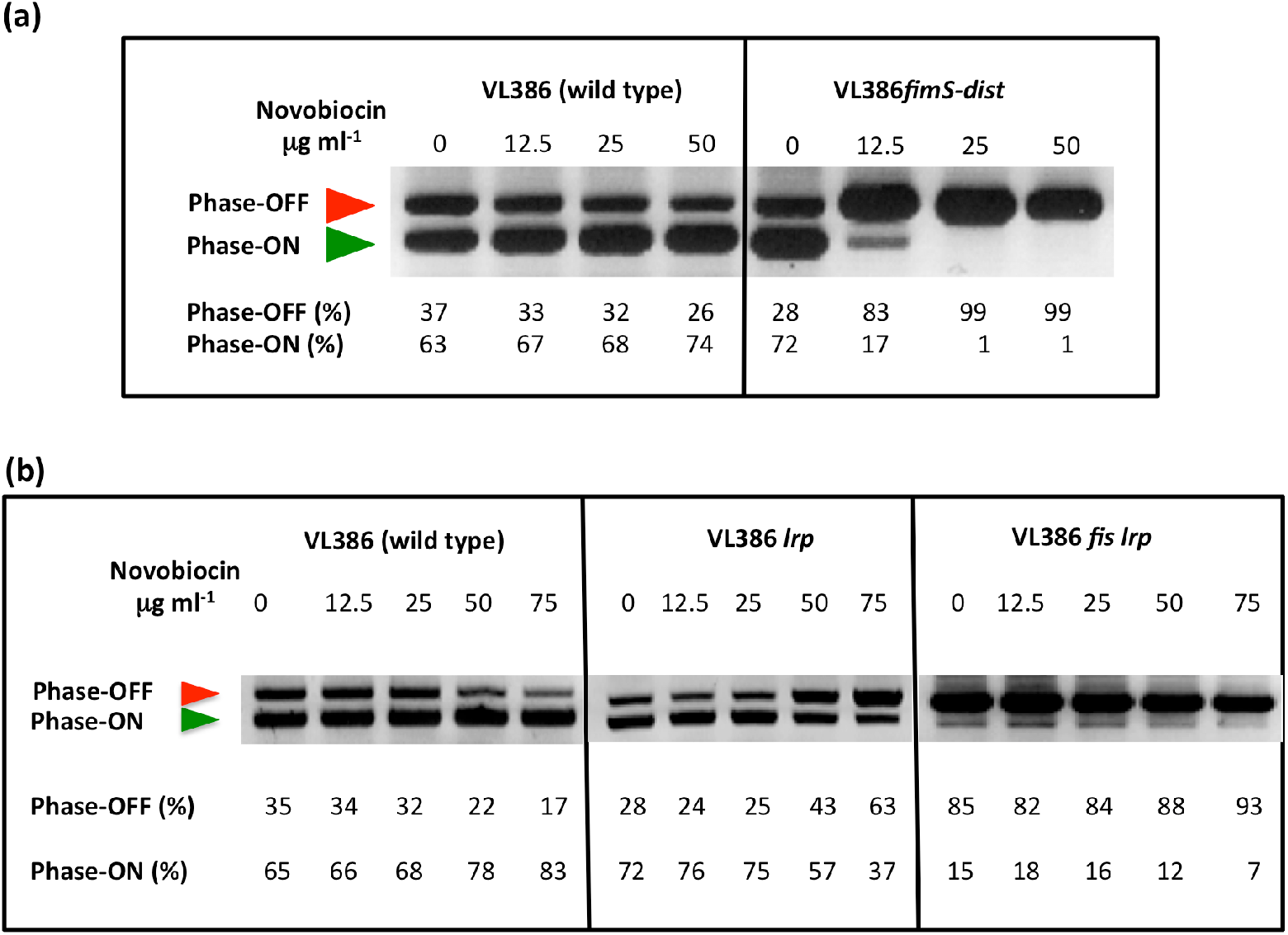
Loss of Fis binding and loss of Fis production biases *fimS* towards Phase-OFF. (a) Electrophoresis of the *fimS* DNA fragments from the wild-type strain (VL386) and from its derivative (VL386*fimS-dist*) in which the Fis binding site in *fimS* is disrupted, following BstUI-digestion of the PCR-amplified *fimS* genetic element. (b) Electrophoresis of the *fimS* DNA fragments from the wild-type strain (VL386), its *lrp* knockout mutant derivative, and the derivative with knockout mutations in both the *lrp* and *fis* genes, following BstUI-digestion of the PCR-amplified *fimS* genetic element. The red arrowhead indicates the 539-bp Phase-OFF diagnostic band and the green arrowhead shows the 433-bp diagnostic band. In (a) and (b), the cultures had been treated with novobiocin at the concentrations given above each gel lane. The intensities of the DNA bands in each lane corresponding to the ON and OFF orientations of *fimS* in each lane were determined by densitometry and are reported as percentages below the lane. The experiment was performed three times and typical data are presented.

### The *fimS* Fis binding site substantially overlaps a binding site for Lrp

The *fimS* DNA sequence that is protected from DNase I digestion by Fis overlaps the previously characterised LRP-2 binding site used by the leucine-responsive regulatory protein, Lrp. This Lrp site helps to determine the inversion bias of *fimS* [Corcoran et al., 2009; Kelly et al., 2006]. We first studied the *fimS* inversion pattern in *lrp* and *lrp fis* knockout mutants with increasing concentrations of novobiocin, compared with the wild-type pattern (Fig. 5b). The wild-type culture followed the usual pattern, with *fimS* becoming progressively biased towards the Phase-ON orientation as the novobiocin concentration increased. The *lrp* mutant became mildly biased towards Phase-OFF, in agreement with previous findings; full Phase-OFF biasing requires the disruption of both LRP-1 and LRP-2 (Kelly et al., 2006). In the *lrp fis* double mutant, the switch was already biased towards Phase-OFF before drug treatment and became almost wholly Phase-OFF as novobiocin was introduced (Fig. 5b).

We next investigated the effect of the base substitutions that abrogated Fis binding to *fimS* on the binding of Lrp to the invertible switch. These sequence changes had affected only 2 bp of the Lrp-protected region at the LRP-2 site (Fig. 6). The 135-bp *fimS* probe used in the Fis EMSA contains both the LRP-1 and LRP-2 sites. Binding of Lrp to *fimS* produces a number of complexes that depend on the occupancy of the LRP-1 and LRP-2 sites, individually and collectively [Gally et al., 1994; Kelly et al., 2006]. Data from EMSA experiments using the *fimS* probe, with and without the Fis binding site mutation, showed that formation of the most electrophoretically retarded *fimS-*Lrp complex was reduced at the highest concentration of purified Lrp (Fig. 6). These data were consistent with the previously described effect of disrupting the LRP-2 site on LRP-DNA complex formation at *fimS* [Kelly et al., 2006] and with the LRP-2 site also being targeted by the Fis protein.

**Fig. 6.**
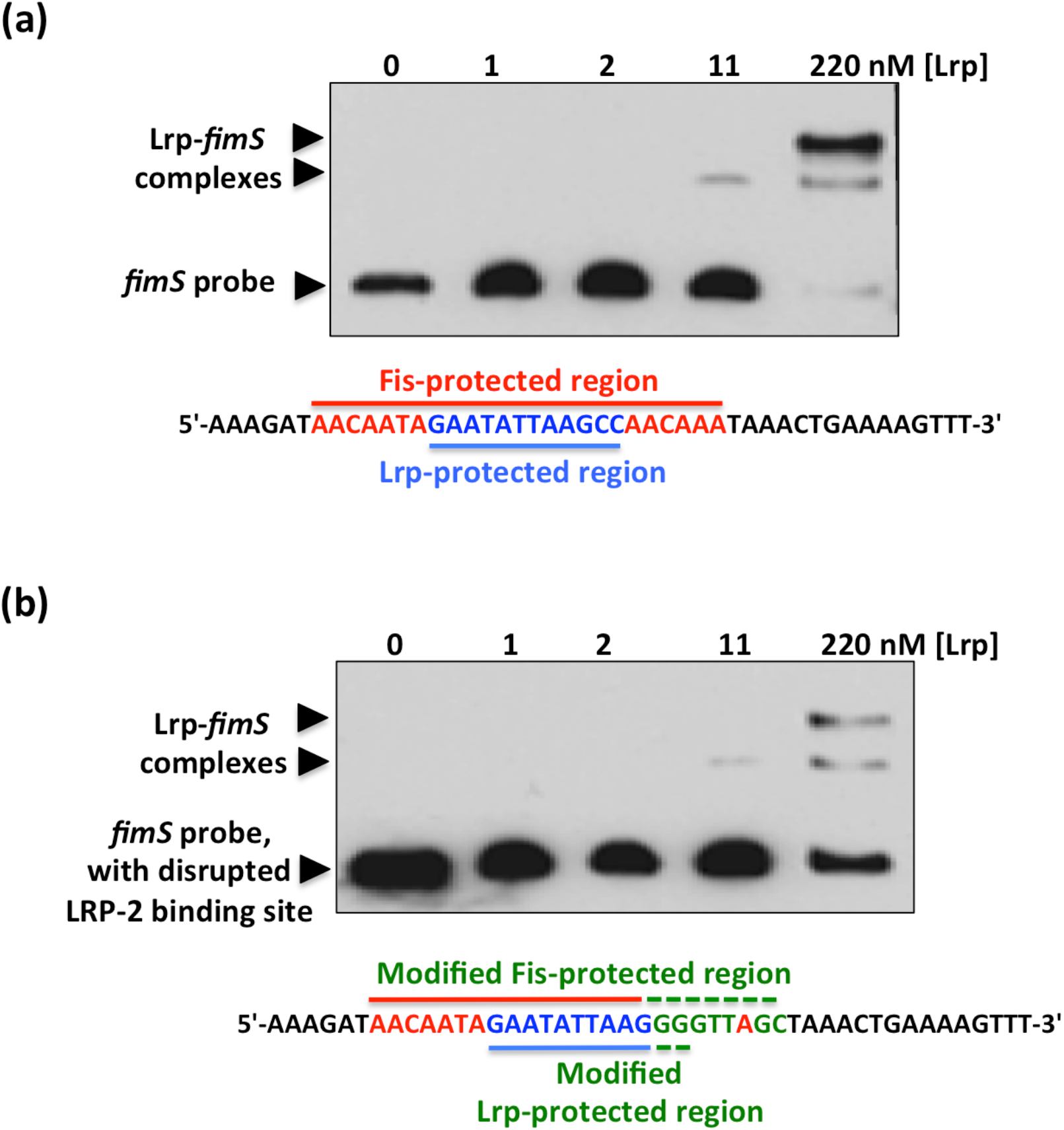
EMSA showing Lrp binding to *fimS* with an intact or a disrupted Fis binding site. (a) Purified Lrp protein was incubated at the concentrations shown with a PCR-generated DNA fragment from *fimS* that contained an intact Fis binding site. The sequence of the region protected from DNase I digestion by Fis is shown in red and the Lrp binding site (LRP-2) is shown in blue. (b) The EMSA was repeated using the *fimS* derivative with the disrupted Fis binding site. The base substitutions are shown below the gel in green, together with the unchanged bases from the Fis binding site (red) and the LRP-2 site (blue). In both (a) and (b), arrowheads show the positions of bands corresponding to the unbound DNA probe and the Lrp-*fimS* complexes.

### Lrp displaces Fis from the *fimS* genetic switch

The data obtained thus far show that the Lrp and Fis proteins both target the LRP-2 binding site in *fimS* (Figs 5a, 5b). Fis is available in high concentration at the onset of exponential growth before becoming rapidly diluted by cell division as the bacteria in the culture expand in numbers. Since Fis and Lrp both influence *fimS* inversion in the same direction, we hypothesised that Fis might be replaced by Lrp at the LRP-2 site when Fis concentration declines as exponential phase progressed. It was also possible that Lrp might actively displace Fis through competition for the same binding site. Therefore, we next assessed the ability of Lrp to displace Fis from LRP-2 in a competitive EMSA. Here, the *fimS* DNA was preloaded with purified Fis at a constant concentration and purified Lrp was added at increasing concentrations (Fig. 7). Lrp and Fis complexes with *fimS* could co-exist at intermediate concentrations of Lrp (22 to 110 nM), presumably indicating occupation of the LRP-1 site by Lrp and of LRP-2 site by Fis, but at the highest concentrations of Lrp, the Fis-*fimS* complex was only weakly detected, presumably because the LRP-2 site was now occupied by Lrp on most *fimS* copies in the reaction (Fig. 7). The EMSA competition showed a specific Fis-*fimS* complex and Lrp-*fimS* complexes; we did not detect evidence of a Fis-plus-Lrp complex with *fimS*, in which Lrp occupied both the LRP-1 and LRP-2 sites with co-binding of Fis and Lrp to LRP-2. Thus, the binding pattern of Fis and Lrp at *fimS* differed from the pattern seen with Fis and Xis at the λ *attR* site, where both Fis and Xis act as directionality determinants in λ excision from the chromosome and both bind to overlapping sites in the DNA simultaneously (Fig. 8) [Seah et al., 2014; Landy, 2015; Papagiannis et al., 2007].

**Fig. 7.**
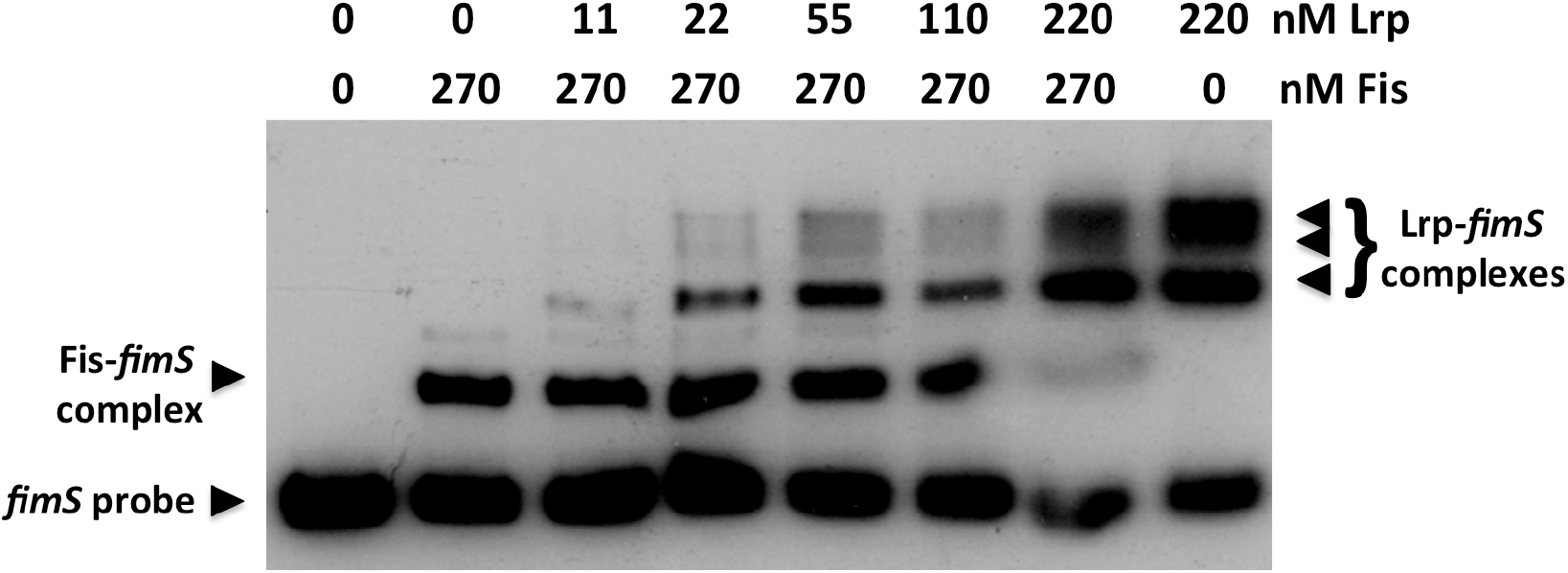
Displacement of Fis from *fimS* by the Lrp protein. Either 0 or 270 nM of purified Fis protein was pre-bound to a fragment of *fimS* DNA containing the LRP-1, LRP-2 binding sites and the Fis binding site. Increasing concentrations of purified Lrp protein were added to the Fis-*fimS* complex and resolved by electrophoresis. Control lanes containing *fimS* DNA with no protein, with just Fis, or with just Lrp were included as controls. Arrowheads indicate the unbound *fimS* probe, the Fis-*fimS* complex, and the Lrp-*fimS* complexes. At 220 nM, Lrp almost completely displaced pre-bound Fis from *fimS*.

**Fig. 8.**
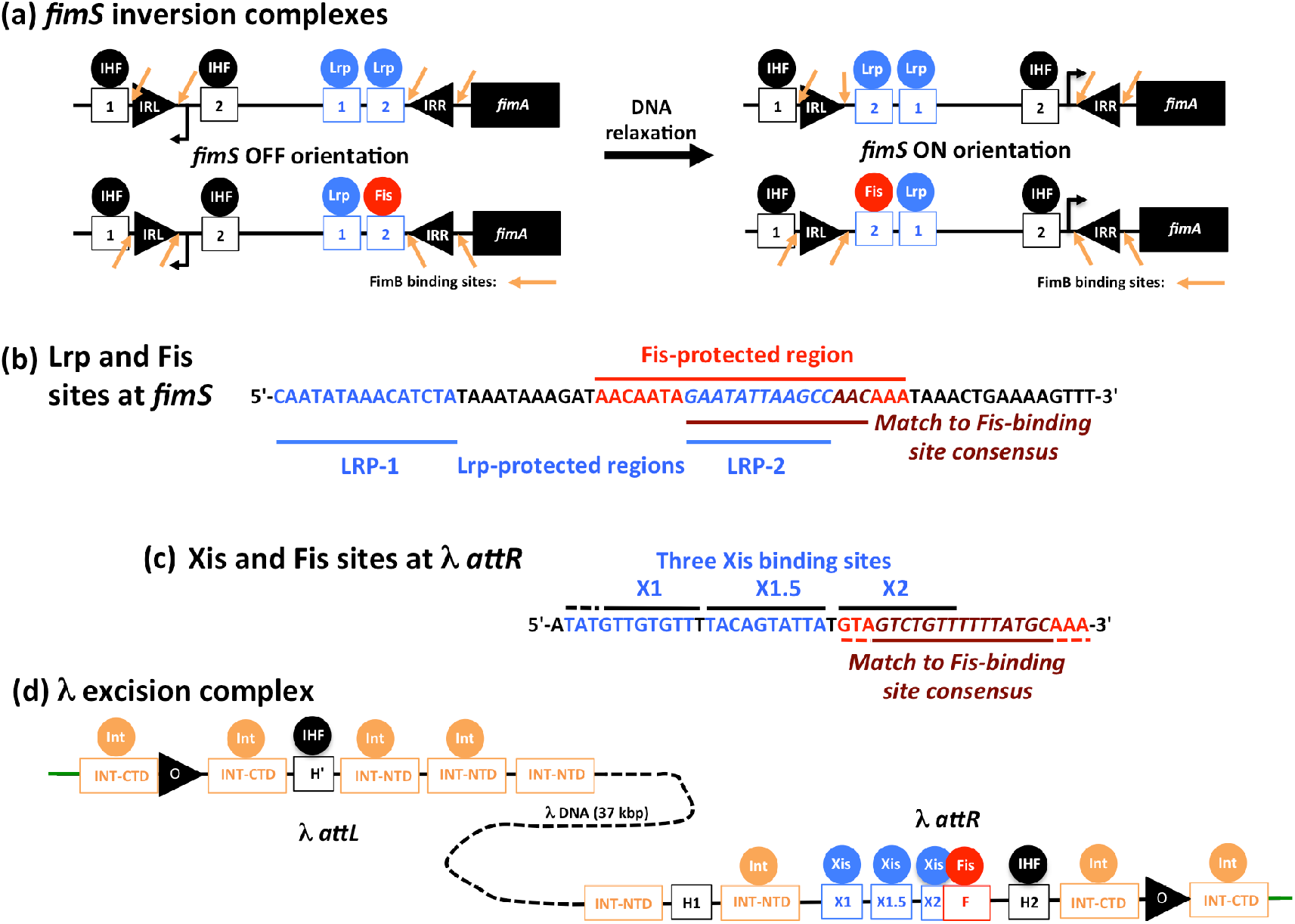
A comparison of the *fimS* inversion complexes and the λ prophage excision complex. (a) The *fimS* genetic switch is shown in the OFF orientation at left (P_*fimA*_ directed away from *fimA*) and in the ON orientation at right (P_*fimA*_ directed towards *fimA*); the angled black arrow represents the P_*fimA*_ promoter. In each complex, both IHF sites are occupied by IHF (black). Alternative states of *fimS* are illustrated, one in which Lrp (blue) occupies both LRP-1 and LRP-2 and one in which LRP-2 is occupied by Fis (red). The horizontal arrow indicates the bias in switching from Phase-OFF to Phase-ON that accompanies DNA relaxation in wild-type strain VL386. Orange arrows show the four binding sites occupied by the FimB recombinase on either side of the IRL and IRR sequences in each complex. Not to scale. (b) A summary of the DNA sequences of the LRP-1 and LRP-2 binding sites (blue) in *fimS*, showing that the LRP-2 (blue italics) site is nested completely within the Fis binding site (red/crimson). (c) The relationships of the three binding sites for the Xis excisionase, X1, X1.5, and X2 (blue) to the binding site for Fis, F (red/crimson) in the λ *attR* region: here, the Fis site overlaps the X2 site completely. (d) The λ prophage excision complex. Bacterial chromosomal DNA is shown in green at the ends of the prophage. Int mediated site-specific recombination takes place within the 7-bp direct repeats represented by the black arrowheads, one each in λ *attL* and *attR*. These mark the boundaries between the prophage and the flanking chromosomal sequences. Int can bind to nine sites (orange rectangles) but in the excision reaction only seven are utilised. INT-NTD and INT-CTD indicate sites of contact for the amino-terminal domain and the carboxyl terminal domain of Int, respectively. IHF (black) occupies only two of its three binding sites; Xis (blue) occupies all three of its sites, and the Fis protein (red) simultaneously occupies its F site, overlapping the Xis binding site X2. Once the prophage is excised, the λ *att* site in the chromosome will consist of just one copy of the direct repeat (black arrowhead) flanked by a single INT-CTD site on either side (not shown). Not to scale.

## DISCUSSION

Our data reveal a delicate interplay between DNA supercoiling/relaxation, LRP, and FIS, in determining the directionality of the FimB-mediated site-specific recombination reaction in *E. coli*. The *fimS* switch becomes progressively biased towards the ON orientation following novobiocin-induced inhibition of DNA gyrase, the topoisomerase that introduces negative supercoils into DNA (Fig. 1b) [Corcoran and Dorman, 2009; Dove and Dorman, 1994; 1996; Kelly et al., 2006; Muller et al., 2009]. In an *lrp* knockout mutant, DNA relaxation results in a reversal of *fimS* inversion outcomes in favour of the OFF, rather than the ON, orientation [Corcoran and Dorman, 2009; Kelly et al., 2006]. Inactivation of Fis production in the *lrp* knockout mutant produces an even stronger preference for the ON orientation, one that is achieved even in the absence of novobiocin, but which becomes much more pronounced as concentrations of the drug increase (Fig. 5). The relationship between Fis and Lrp at *fimS* has similarities to the relationship between Fis and the Xis directionality determinant at *attR* in bacteriophage λ excision (Fig. 8).

Int-mediated excisive recombination of bacteriophage λ is enhanced when DNA supercoiling levels are low, whereas integrative recombination requires negative supercoiling of the phage DNA [Crisona et al., 1991; Seah et al., 2014]. The phage-encoded Xis architectural protein stimulates excision by a factor of 10^6^ while simultaneously inhibiting reintegration of the phage [Abremski and Gottesman, 1982; Nash, 1975]. Xis binds to the X1, X1.5, and X2 sites in the *attR* arm of the λ prophage to form a microfilament [Abbani et al., 2007], with site X2 overlapping a binding site for Fis, the F site [Thompson et al., 1987] (Fig. 8). Initially it was thought that Fis substituted for Xis at site X2, allowing excision to proceed under conditions where Xis was limiting [Landy, 2015]. It is now understood that Xis occupies all three of its binding sites, with Fis binding simultaneously to its F site (Fig. 8), producing a nucleoprotein complex with a DNA conformation that is optimal for excisive recombination [Abbani et al., 2007; Papagiannis et al., 2007]. The role Fis plays in recruiting Xis does not seem to involve protein-protein contact, but is achieved through DNA allostery [Hancock et al., 2019]. While Xis imposes a preference for excision on the λ prophage, the Fis protein has been reported to stimulate integration as well as excision [Esposito and Gerard, 2003], especially in the absence of Xis [Ball and Johnson, 1991].

Thus, the λ excision complex differs from the *fimS* OFF to ON inversion complex in that Xis and Fis bind together to overlapping sites in *attR*, while Lrp and Fis bind competitively to overlapping sites in *fimS* (Fig.7). Despite this distinction, the two systems share a preference for relaxed DNA to facilitate a direction-specific recombination reaction; a dependency on tyrosine-integrase recombinases to catalyse the reaction; and a requirement for IHF to occupy two sites in the DNA to organise a recombination substrate with an appropriate architecture (Figs 1a, 8).

Int differs from the FimB and FimE recombinases in that it makes contact with the DNA through both its amino-terminal and its carboxyl-terminal domains (NTD and CTD, respectively) at up to nine sites [Landy, 2015]. The smaller fimbrial recombinases lack the corresponding NTD and make contact with *fimS* at four sites only, two flanking each of the 9-bp inverted repeats, IRL and IRR, that form the boundaries of *fimS* [Kulasekara and Blomfield, 1999; McCusker et al., 2008] (Fig. 8). DNA cleavage and ligation takes place within these inverted *fimS* repeats [McCusker et al., 2008] while Int cleaves and religates the λ prophage within the 7-bp inverted repeats that are flanked by Int-CTD binding sites in *attL* and *attR* [Landy, 2015]. In λ site-specific recombination, Fis can stimulate both integration and excision; in *fimS*, Fis plays a role as a directionality co-determinant with Lrp, favouring the ON-to-OFF reaction.

DNA relaxation is a feature of stationary phase cultures [Bordes et al., 2003; Conter et al., 1997] and is a condition that biases *fimS* towards the ON phase [Corcoran and Dorman, 2009; Dove and Dorman, 1994; 1996; Kelly et al., 2006; Muller et al., 2009]. This bias requires both Fis and Lrp (Figs. 5b). Loss of either protein introduces an alternative bias towards the OFF phase as DNA relaxes [Corcoran et al., 2009; Kelly et al., 2006]; loss of both proteins results in *fimS* being maintained in the OFF phase in almost all bacteria in the population (Fig. 5b).

The competitive relationship described here for Fis and Lrp at *fimS* is reminiscent of the competition between the Dam methylase and Lrp for access to overlapping sites in the regulatory region of *pap*, the operon that encodes Pap pili in uropathogenic strains of *E. coli* [Hernday et al., 2003; Peterson and Reich, 2008]. The *fim and pap* operons engage in regulatory crosstalk via PapB-mediated repression of *fim* operon transcription [Holden et al., 2001; Lindberg et al., 2008; Xia et al., 2000]. Although DNA recombination does not contribute to the operation of the phase-variable *pap* switch, the outcome of the Dam/Lrp competition determines whether the *pap* operon will or will not be transcribed. Lrp accumulates in stationary phase cultures growing in rich media [Landgraf et al., 1996] while Dam concentrations decline under those same growth conditions [Rasmussen et al., 1994]. It has been suggested that the shift in the Dam/Lrp balance in favour of Lrp facilitates a shift in Pap production from the ON to the OFF phase [Peterson and Reich, 2008]. These features of *pap* gene regulation by Dam and Lrp mirror those described here for *fim* gene regulation by Fis and Lrp. Like Dam, Fis is produced in decreasing amounts as stationary phase approaches, while the production of Lrp increases [Landgraf et al., 1996]. In early exponential phase, the abundant Fis protein collaborates with the less abundant Lrp to bias *fimS* towards the OFF phase, expanding the number of afimbriate, planktonic bacteria in the population. As exponential growth gives way to growth stasis, the Fis concentration declines sharply and Lrp replaces it at the LRP-2 site in *fimS*, if necessary by competitive displacement.

Biasing the *fimS* switch towards the OFF orientation when Fis is abundant and DNA is negatively supercoiled, links *fim* switch inversion preferences to bacterial physiology. The Fis protein is maximally abundant at the beginning of the exponential phase of growth and is almost undetectable by the onset of stationary phase [Ball et al., 1992; Bogue et al., 2020; O Cróinín and Dorman 2007; Osuna et al., 1995]. As Fis levels decline, Lrp is available to replace it at *fimS*, prolonging the bias towards Phase-OFF as long as the DNA remains negatively supercoiled. However, stationary phase is a period of reduced metabolic flux, producing a reduced [ATP]/[ADP] ratio that is unfavourable for the negative DNA supercoiling activity of DNA gyrase [Conter et al., 1997; Hsieh et al., 1991a; 1991b; van Workum et al., 1996; Westerhoff and van Workum, 1990]. At this stage of the growth cycle, DNA becomes relaxed [Bordes et al., 2003; Conter et al., 1997], a condition that biases the *fimS* switch to the ON phase in the presence of Lrp [Corcoran and Dorman, 2009; Dove and Dorman, 1994; 1996; Kelly et al., 2006; Muller et al., 2009]. Removal of Lrp from *fimS* reverses the inversion bias back towards the OFF phase [Kelly et al., 2006].

The increased representation of fimbriate bacteria in the population of late-exponential phase/stationary phase cells promotes bacterial attachment to abiotic and biological surfaces, with an associated production of biofilm [Muller et al., 2009; Pratt and Kolter, 1998; Schembri et al., 2003]. The resulting transition from a planktonic to a community based, attached lifestyle within the protective shield of a biofilm, enhances the survival chances of the bacterial population during a period of unfavourable environmental conditions. Overall, our findings describe a molecular mechanism by which the bet-hedging strategy represented by the stochastic inversion of *fimS* is suspended in favour of the more deterministic outcome of ensuring that type 1 fimbriae are produced by a majority of bacteria in the population. Fis and Lrp are required, in association with DNA relaxation, for the implementation of this deterministic strategy.

## Acknowledgements

We are grateful to Stephen GJ Smith for insightful discussions.

## Author contributions

Conceptualisation: CJD; Data analysis: CC; MCB; CJD; Funding acquisition: CJD; Investigation: CC; MCB; CJD; Methodology: CC; Project administration: CJD; Resources: CJD; Supervision: CJD; Writing – original draft: CJD; Writing – review and editing: CC; MCB; CJD.

## References

Abbani MA, Papagiannis CV, Sam MD, Cascio D, Johnson RC, et al. Structure of the cooperative excisionase (Xis)-DNA complex reveals a micronucleoprotein filament that regulates phage lambda intasome assembly. Proc Natl Acad Sci USA 2007;104:2109–2114.

Abraham JM, Freitag CS, Clements JR, Eisenstein BI. An invertible element of DNA controls phase variation of type 1 fimbriae of *Escherichia coli*. Proc Natl Acad Sci USA 1985;82:5724–5727.

Abremski K, Gottesman S. Purification of the bacteriophage λ *xis* gene product required for λ excisive recombination. J Biol Chem 1982;257:9658–9662.

Anderson GG, Palermo JJ, Schilling JD, Roth R, Heuser J, et al. Intracellular bacterial biofilm-like pods in urinary tract infections. Science 2003;301:105–107.

Ball CA, Johnson RC. Multiple effects of Fis on integration and the control of lysogeny in phage λ. J Bacteriol 1991;173(13):4032–4038.

Ball CA, Osuna R, Ferguson KC, Johnson RC. Dramatic changes in Fis levels upon nutrient upshift in *Escherichia coli*. J Bacteriol 1992;174: 8043–8056.

Bétermier M, Lefrère V, Koch C, Alazard R, Chandler M. The *Escherichia coli* protein, Fis: specific binding to the ends of phage Mu DNA and modulation of phage growth. Mol Microbiol 1989;3(4):459–468.

Blomfield IC, McClain MS, Princ JA, Calie PJ, Eisenstein BI. Type 1 fimbriation and *fimE* mutants of *Escherichia coli* K-12. J Bacteriol 1991;173(17):5298–5307.

Blomfield IC, Vaughn V, Rest RF, Eisenstein BI. Allelic exchange in *Escherichia coli* using the *Bacillus subtilis sacB* gene and a temperature-sensitive pSC101 replicon. Mol Microbiol 1991b;5:1447–1457.

Blomfield IC, Calie PJ, Eberhardt KJ, McClain MS, Eisenstein BI. Lrp stimulates phase variation of type 1 fimbriation in *Escherichia coli* K-12. J Bacteriol 1993;175(1):27–36.

Blomfield IC, Kulasekara DH, Eisenstein BI. Integration host factor stimulates both FimB- and FimE-mediated site-specific DNA inversion that controls phase variation of type 1 fimbriae expression in *Escherichia coli*. Mol Microbiol 1997;23(4):705–117.

Bogue MM, Mogre A, Beckett, MC, Thomson NR, Dorman CJ. Network rewiring: physiological consequences of reciprocally exchanging the physical locations and growth-phase-dependent expression patterns of the *Salmonella fis* and *dps* genes. mBio 2020;11(5):e0218–20.

Bordes P, Conter A, Morales V, Bouvier J, Kolb A, Gutierrez C. DNA supercoiling contributes to disconnect sigmaS accumulation from sigmaS-dependent transcription in *Escherichia coli*. Mol Microbiol 2003;48:561–571.

Calvo JM, Matthews RG. The Leucine-responsive Regulatory Protein, a global regulator of metabolism in *Escherichia coli*. Microbiol Rev 1994;58(3):466–490.

Cameron AD, Stoebel DM, Dorman CJ. DNA supercoiling is differentially regulated by environmental factors and Fis in *Escherichia coli* and *Salmonella enterica*. Mol Microbiol 2011;80(1):85–101.

Chen S, Calvo JM. Leucine-induced dissociation of *Escherichia coli* Lrp hexadecamers to octamers. J Mol Biol 2002;318:1031–1042.

Chen S, Iannolo M, Calvo JM. Cooperative binding of the leucine-responsive regulatory protein (Lrp) to DNA. J Mol Biol 2005;345:251–264.

Cho B-K, Barrett CL, Knight EM, Park YS, Palsson BØ. Genome-scale reconstruction of the Lrp regulatory network in *Escherichia coli*. Proc Natl Acad Sci USA 2008;105:19462–19467.

Connell I, Agace W, Klemm P, Schembri M, Mårild S, Svanborg C. Type 1 fimbrial expression enhances *Escherichia coli* virulence for the urinary tract. Proc Natl Acad Sci USA 1996;93: 9827–9832.

Conter A, Menchon C, Gutierrez C. Role of DNA supercoiling and *rpoS* sigma factor in the osmotic and growth phase-dependent induction of the gene *osmE* of *Escherichia coli* K12. J Mol Biol 1997;273:75–83.

Corcoran CP, Dorman CJ. DNA relaxation-dependent phase biasing of the *fim* genetic switch in *Escherichia coli* depends on the interplay of H-NS, IHF and LRP. Mol Microbiol 2009;74(5):1071–1082.

Crisona NJ, Weinberg RL, Peter BJ, Sumners DW, Cozzarelli NR. The topological mechanism of phage λ integrase. J Mol Biol 1999;289(4):747–775.

Cui Y, Wang Q, Stormo GD, Calvo JM. A consensus sequence for binding of Lrp to DNA. J Bacteriol 1995;177:4872–4880.

Dorman CJ, Higgins CF. Fimbrial phase variation in *Escherichia coli*: dependence on integration host factor and homologies with other site-specific recombinases. J Bacteriol 1987;169(8): 3840–3843.

Dove SL, Dorman CJ. The site-specific recombination system regulating expression of the type 1 fimbrial subunit gene of *Escherichia coli* is sensitive to changes in DNA supercoiling. Mol Microbiol 1994;14:975–988.

Dove SL, Dorman CJ. Multicopy *fimB* gene expression in *Escherichia coli*: binding to inverted repeats in vivo, effect on *fimA* gene transcription and DNA inversion. Mol Microbiol 1996;21:1161–1173.

Eisenstein BI. Phase variation of type 1 fimbriae in *Escherichia coli* is under transcriptional control. Science 1981;214:347–349.

Eisenstein BI, Sweet DS, Vaughn V, Friedman DI. Integration host factor is required for the DNA inversion that controls phase variation in *Escherichia coli*. Proc Natl Acad Sci USA 1987;84(18):6506–6510.

Esposito D, Gerard GF. The *Escherichia coli* Fis protein stimulates bacteriophage λ integrative recombination in vitro. J Bacteriol 2003;185(10):3076–3080.

Flores-Mireles AL, Walker JN, Caparon MG, Hultgren SJ. Urinary tract infections: epidemiology, mechanisms of infection and treatment options. Nat Rev Microbiol 2015;13:269–284.

Gally DL, Rucker TJ, Blomfield IC. The Leucine-responsive Regulatory Protein binds to the *fim* switch to control phase variation of type 1 fimbrial expression in *Escherichia coli* K-12. Mol Microbiol 1994;176(18):5665–5672.

Gally DL, Leathart J, Blomfield IC. Interaction of FimB and FimE with the *fim* switch that controls the phase variation of type 1 fimbriae in *Escherichia coli* K-12. Mol Microbiol 1996;21(4):725–738.

Gille H, Egan JB, Roth A, Messer W. The Fis protein binds and bends the origin of chromosomal DNA replication, *oriC*, of *Escherichia coli*. Nucleic Acids Res 1991;19(15):4167–4172.

Gosink KK, Ross W, Leirmo S, Osuna R, Finkel SE, Johnson RC, et al. DNA binding and bending are necessary but not sufficient for Fis-dependent activation of *rrnB* P1. J Bacteriol 1993;175(6):1580–1589.

Grimwade JE, Leonard AC. Blocking, bending, and binding: regulation of initiation of chromosome replication during the *Escherichia coli* cell cycle by transcriptional modulators that interact with origin DNA. Front Microbiol 2021;12:732270.

Hanahan D. Studies on transformation of *Escherichia coli* with plasmids. J Mol Biol 1983;166:557–580.

Hancock SP, Stella S, Cascio D, Johnson RC. DNA sequence determinants controlling affinity, stability and shape of DNA complexes bound by the nucleoid protein Fis. PLoS One 2016;11(3):e0150189.

Hancock SP, Cascio D, Johnson RC. Cooperative DNA binding by proteins through DNA shape complementarity. Nucleic Acids Res 2019;47(16):8874–8887.

Hernday AD, Braaten BA, Low DA. The mechanism by which DNA adenine methylase and PapI activate the *pap* epigenetic switch. Mol Cell 2003;12(4):947–957.

Higgins CF, Dorman CJ, Stirling DA, Waddell L, Booth IR, May G, et al. A physiological role for DNA supercoiling in the osmotic regulation of gene expression in *S. typhimurium* and *E. coli*. Cell 1988;52(4):569–584.

Hinde P, Deighan P, Dorman CJ. Characterization of the detachable Rho-dependent transcription terminator of the *fimE* gene in *Escherichia coli* K-12. J Bacteriol 2005;187(24):8256–8266.

Holden NJ, Uhlin BE, Gally DL. PapB paralogues and their effect on the phase variation of type 1 fimbriae in *Escherichia coli*. Mol Microbiol 2001;42:319–330.

Holden NJ, Blomfield IC, Uhlin BE, Totsika M, Kulasekara DH, Gally DL. Comparative analysis of FimB and FimE recombinase activity. Microbiology 2007;153(Pt 12):4138–4149.

Hsieh LS, Burger RM, Drlica K. Bacterial DNA supercoiling and [ATP]/[ADP]. Changes associated with a transition to anaerobic growth. J Mol Biol 1991;219:443–450.

Hsieh LS, Rouvière-Yaniv J, Drlica K. Bacterial DNA supercoiling and [ATP]/[ADP] ratio: changes associated with salt shock. J Bacteriol 1991;173:3914–3917.

Inoue Y, Tanaka H, Kasho K, Fujimitsu K, Oshima T, et al. Chromosomal location of the DnaA-reactivating sequence DARS2 is important to regulate timely initiation of DNA replication in *Escherichia coli*. Genes Cells 2016;21(9):1015–1023.

Jiang X, Brantley Hall A, Arthur TD, Plichta DR, Covington CT, et al. Invertible promoters mediate bacterial phase variation, antibiotic resistance, and host adaptation in the gut. Science 2019;363(6423):181–187.

Joyce SA, Dorman CJ. A Rho-dependent phase-variable transcription terminator controls expression of the FimE recombinase in *Escherichia coli*. Mol Microbiol 2002;45(4):1107–1117.

Justice SS, Hung C, Theriot JA, Fletcher DA, Anderson GG, et al. Differentiation and developmental pathways of uropathogenic *Escherichia coli* in urinary tract pathogenesis. Proc Natl Acad Sci USA 2004;101:1333–1338.

Karambelkar S, Swapna G, Nagaraja V. Silencing of toxic gene expression by Fis. Nucleic Acids Res 2012;40(10):4358–4367.

Keane OM, Dorman CJ. The *gyr* genes of *Salmonella enterica* serovar Typhimurium are repressed by the factor for inversion stimulation, Fis. Mol Genet Genomics 2003;270(1):56–65.

Kelly A, Goldberg MD, Carroll RK, Danino V, Hinton JCD, Dorman CJ. A global role for Fis in the transcriptional control of metabolic and type III secretion genes of *Salmonella enterica* serovar Typhimurium. Microbiology 2004;150(7):2037–2053.

Kelly A, Conway C, O Cróinín T, Smith SGJ, Dorman CJ. DNA supercoiling and the Lrp protein determine the directionality of *fim* switch DNA inversion in *Escherichia coli* K-12. J Bacteriol 2006;188(15):5356–5363.

Klemm, P. Two regulatory *fim* genes, *fimB* and *fimE*, control the phase variation of type 1 fimbriae in *Escherichia coli*. EMBO J 1986;5:1389–1393.

Kroner GM, Wolfe MB, Freddolino PL. *Escherichia coli* Lrp regulates one-third of the genome via direct, cooperative, and indirect routes. J Bacteriol 2019;201(3):e00411–18.

Kulasekara HD, Blomfield IC. The molecular basis for the specificity of *fimE* in the phase variation of type 1 fimbriae of *Escherichia coli* K-12. Mol Microbiol 1999;31(4):1171–1181.

Landgraf JR, Wu J, Calvo JM. Effects of nutrition and growth rate on Lrp levels in *Escherichia coli*. J Bacteriol 1996;178:6930–6936.

Landy A. The λ integrase site-specific recombination pathway. Microbiol Spectr 2015; 3(2):MDNA3-0051-2014.

Lindberg S, Xia Y, Sonden B, Göransson M, Hacker J, et al. Regulatory interactions among adhesin gene systems of uropathogenic *Escherichia coli*.. Infect Immun 2008;76:771–780.

McClain MS, Blomfield IC, Eisenstein BI. Roles of *fimB* and *fimE* in site-specific DNA inversion associated with phase variation of type 1 fimbriae in *Escherichia coli*. J Bacteriol 1991;173: 5308–5314.

McCusker MP, EC Turner, CJ Dorman. DNA sequence heterogeneity in site-specific recombinase binding sites determines the directionality of *fim* genetic switch DNA inversion in *Escherichia coli* K-12. Mol Microbiol 2008;67(1):171–187.

McFarland KA, Dorman CJ. Autoregulated expression of the gene coding for the leucine-responsive protein, Lrp, in *Salmonella enterica* serovar Typhimurium. Microbiology 2008;154(7):2008–2016.

McFarland KA, Lucchini S, Hinton JCD, Dorman CJ. The Leucine-responsive Regulatory Protein, Lrp, activates transcription of the *fim* operon in *Salmonella enterica* Serovar Typhimurium via the *fimZ* regulatory gene. J Bacteriol 2008; 190(2):602–612.

Miller JH. A Short Course in Bacterial Genetics. Cold Spring Harbor Laboratory Press, 1992; Cold Spring Harbor, New York.

Muller CM, Aberg A, Straseviciene J, Emody L, Uhlin BE, et al. Type 1 fimbriae, a colonization factor of uropathogenic *Escherichia coli*, are controlled by the metabolic sensor CRP-cAMP. PLoS Pathog 2009;5(2):e1000303.

Muskhelishvili G, Travers A. Transcription factor as a topological homeostat. Front Biosci 2003;8:d279–285.

Nakano Y, Yoshida Y, Yamashita Y, Koga T. Construction of a series of pACYC-derived plasmid vectors. Gene 1995;162:157–158.

Nash HA. Integrative recombination of bacteriophage lambda DNA in vitro. Proc Natl Acad Sci USA 1975;72:1072–1076.

Ó Cróinín T, Dorman CJ. Expression of the Fis protein is sustained in late-exponential and stationary-phase cultures of *Salmonella enterica* serovar Typhimurium grown in the absence of aeration. Mol Microbiol 2007;66:237–251.

Ó Cróinín T, Carroll RK, Kelly A, Dorman CJ. Roles for DNA supercoiling and the Fis protein in modulating expression of virulence genes during intracellular growth of *Salmonella enterica* serovar Typhimurium. Mol Microbiol 2006;62(3):869–882.

O’Gara JP, Dorman CJ. Effects of local transcription and H-NS on inversion of the *fim* switch of *Escherichia coli*. Mol Microbiol 2000;36(2):457–466.

Kawula TH, Orndorff PE. Rapid site-specific DNA inversion in *Escherichia coli* mutants lacking the histonelike protein H-NS. J Bacteriol 1991;173(13):4116–4123.

Osuna R, Lienau D, Hughes KT, Johnson RC. Sequence, regulation, and functions of *fis* in *Salmonella typhimurium*. J Bacteriol 1995;177:2021–2032.

Papagiannis CV, Sam MD, Abbani MA, Yoo D, Cascio D, et al. Fis targets assembly of the Xis nucleoprotein filament to promote excisive recombination by phage lambda. J Mol Biol 2007;367:328–343.

Peterson SN, Reich NO. Competitive Lrp and Dam assembly at the *pap* regulatory region: implications for mechanisms of epigenetic regulation. J Mol Biol 2008;383(1):92–105.

Pratt LA, Kolter R. Genetic analysis of *Escherichia coli* biofilm formation: roles of flagella, motility, chemotaxis and type I pili. Mol Microbiol 1998;30:285–293.

Rasmussen LJ, Marinus MG, Lobner-Olesen A. Novel growth rate control of *dam* gene expression in *Escherichia coli*. Mol Microbiol 1994;12:631–638.

Rochman M, Aviv M, Glaser G, Muskhelishvili G. Promoter protection by a transcription factor acting as a local topological homeostat. EMBO Rep 2002;3(4):355–360.

Ryan VT, Grimwade JE, Camara JE, Crooke E, Leonard AC. *Escherichia coli* prereplication complex assembly is regulated by dynamic interplay among Fis, IHF and DnaA. Mol Microbiol 2004;51(5):1347–1359.

Saldaña-Ahuactzi Z, Soria-Bustos J, Martínez-Santos VI, Yañez-Santos JA, Martínez-Laguna Y, et al. The Fis nucleoid protein negatively regulates the phase variation *fimS* switch of the type 1 pilus operon in Enteropathogenic *Escherichia coli*. Front Microbiol 2022;13:882563.

Sambrook J, Russell D. (2001) Molecular Cloning: A Laboratory Guide. Cold Spring Harbor Laboratory Press, 2001; Cold Spring Harbor, New York.

Sánchez-Romero MA, Casadesús J. Waddington’s landscapes in the bacterial world. Front Microbiol 2021;12:685080.

Schembri MA, Hjerrild L, Gjermansen M, Klemm P. Differential expression of the *Escherichia coli* autoaggregation factor antigen 43. J Bacteriol 2003;185:2236–2242.

Schneider R, Travers A, Kutateladze T, Muskhelishvili G. A DNA architectural protein couples cellular physiology and DNA topology in *Escherichia coli*. Mol Microbiol 1999;34(5):953–964.

Schneider R, Travers A, Muskhelishvili G. The expression of the *Escherichia coli fis* gene is strongly dependent on the superhelical density of DNA. Mol Microbiol 2000;38(1):167–175.

Seah N, Tong W, Warren D, Van Duyne GD, Landy A. Nucleoprotein architectures regulating the directionality of viral integration and excision. Proc Natl Acad Sci USA 2014;111:12372–12377.

Smith SGJ, Dorman CJ. Functional analysis of the FimE integrase of *Escherichia coli* K-12: isolation of mutant derivatives with altered DNA inversion preferences. Mol Microbiol 1999;34(5):965–979.

Spaulding CN, Schreiber HL, Zheng W, Dodson KW, Hazen JE, et al. Functional role of the type 1 pilus rod structure in mediating host-pathogen interactions. eLife 2018;7:e31662.

Stella S, Cascio D, Johnson RC. The shape of the DNA minor groove directs binding by the DNA-bending protein Fis. Genes Dev 2010;15;24(8):814–826.

Sternberg NL, Maurer R. Bacteriophage-mediated generalized transduction in *Escherichia coli* and *Salmonella typhimurium*. Methods Enzymol 1991;204:18–43.

Tani TH, Khodursky A, Blumenthal RM, Brown PO, Matthews RG. Adaptation to famine: A family of stationary-phase genes revealed by microarray analysis. Proc Natl Acad Sci USA 2002;99:13471–13476.

Thompson JF, Moitoso de Vargas L, Koch C, Kahmann R, Landy A. Cellular factors couple recombination with growth phase: characterization of a new component in the lambda site-specific recombination pathway. Cell 1987;50(6):901–908.

van Workum M, van Dooren SJ, Oldenburg N, Molenaar D, Jensen PR, Snoep JL, et al. DNA supercoiling depends on the phosphorylation potential in *Escherichia coli*. Mol Microbiol 1996;20:351–360.

Wang Q, Wu J, Friedberg D, Platko J, Calvo JM. Regulation of the *Escherichia coli lrp* gene. J Bacteriol 1994;176:1831–1839.

Weinreich MD, Reznikoff WS. Fis plays a role in Tn*5* and IS*50* transposition. J Bacteriol 1992;174(14):4530–4537.

Weinstein-Fischer D, Altuvia S. Differential regulation of *Escherichia coli* topoisomerase I by Fis. Mol Microbiol 2007;63(4):1131–1144.

Welch RA, Burland V, Plunkett G, Redford P, Roesch P, et al. Extensive mosaic structure revealed by the complete genome sequence of uropathogenic *Escherichia coli*. Proc Natl Acad Sci USA 2002;99:17020–17024.

Westerhoff HV, van Workum M. Control of DNA structure and gene expression. Biomed Biochim Acta 1990;49:839–853.

Wilson RL, Libby SJ, Freet AM, Boddicker JD, Fahlen TF, et al. Fis, a DNA nucleoid-associated protein, is involved in *Salmonella typhimurium* SP-1 invasion gene expression. Mol Microbiol 2001;22:327–338.

Wright KJ, Seed PC, Hultgren SJ. Development of intracellular bacterial communities of uropathogenic *Escherichia coli* depends on type 1 pili. Cell Microbiol 2007;9:2230–2241.

Xia Y, Gally D, Forsman-Semb K, Uhlin BE. Regulatory cross-talk between adhesin operons in *Escherichia coli*: inhibition of type 1 fimbriae expression by the PapB protein. EMBO J 2000;19:1450–1457.

Xing J, Marshall JS. Mechanics of biofilms formed of bacteria with fimbriae appendages. PLoS One 2020;15(12): e0243280.

Yanisch-Perron C, Viera J, Messing J. Improved M13 phage cloning vectors and host strains: nucleotide sequences of the M13mp18 and pUC19 vectors. Gene 1985;33:103–119.

Zamora M, Ziegler CA, Freddolino PL, Wolfe AJ. A thermosensitive, phase-variable epigenetic switch: *pap* revisited. Microbiol Mol Biol Rev 2020;84(3):e00030–17.

Ziegler CA, Freddolino PL. The leucine-responsive regulatory proteins/feast-famine regulatory proteins: an ancient and complex class of transcriptional regulators in bacteria and archaea. Crit Rev Biochem Mol Biol 2021;56(4):373–400.

Zinser ER, Kolter R. Prolonged stationary-phase incubation selects for *lrp* mutations in *Escherichia coli* K-12. J Bacteriol 2000;182:4361–4365.

